# Alpha-synuclein preformed fibril-induced aggregation and dopaminergic cell death in cathepsin D overexpression and ZKSCAN3 knockout mice

**DOI:** 10.1101/2024.09.18.613763

**Authors:** Toni Mueller, Parker Jeffrey, Yecheng He, Xiaosen Ouyang, David Westbrook, Victor Darley-Usmar, Matthew S. Goldberg, Laura Volpicelli-Daley, Jianhua Zhang

## Abstract

α-synuclein accumulation is recognized as a prominent feature in the majority of Parkinson’s disease cases and also occurs in a broad range of neurodegenerative disorders including Alzheimer’s disease. It has been shown that α-synuclein can spread from a donor cell to neighboring cells and thus propagate cellular damage, antagonizing the effectiveness of therapies such as transplantation of fetal or iPSC derived dopaminergic cells. As we and others previously have shown, insufficient lysosomal function due to genetic mutations or targeted disruption of cathepsin D can cause α-synuclein accumulation. We here investigated whether overexpression of cathepsin D or knockout (KO) of the transcriptional suppressor of lysosomal biogenesis ZKSCAN3 can attenuate propagation of α-synuclein aggregation and cell death. We examined dopaminergic neurodegeneration in the substantia nigra using stereology of tyrosine hydroxylase-immunoreactive cells 4 months and 6 months after intrastriatal injection of α-synuclein preformed fibrils or monomeric α-synuclein control in control, central nervous system (CNS)-cathepsin D overexpressing and CNS-specific ZKSCAN3 KO mice. We also examined pS129-α-synuclein aggregates in the substantia nigra, cortex, amygdala and striatum. The extent of dopaminergic neurodegeneration and pS129-α-synuclein aggregation in the brains of CNS-specific ZKSCAN3 knockout mice and CNS-cathepsin D overexpressing mice was similar to that observed in wild-type mice. Our results indicate that neither enhancing cathepsin D expression nor disrupting ZKSCAN3 in the CNS is sufficient to attenuate pS129-α-synuclein aggregate accumulation or dopaminergic neurodegeneration.

## Introduction

Parkinson’s disease affects approximately 1 in 100 Americans older than 60, and it is estimated that between 2010 and 2030, the number of individuals 65 years or older with Parkinson’s disease will increase by 77% (1, 2). World-wide, Parkinson’s disease affects 1-3% of the population age 50 years or older (3). There are currently no available pharmacological approaches for preventing or attenuating neuronal loss in Parkinson’s disease; consequently, treatment options are limited to symptom management.

α-Synuclein aggregation is recognized as a prominent neuropathological feature of Parkinson’s disease (4) and also occurs in a broad range of neurodegenerative disorders including Alzheimer’s disease (5, 6). α-Synuclein overabundance has been shown to cause cytotoxicity in cell and animal models (7, 8). α-Synuclein is a 140 amino acid protein expressed in neurons. α-Synuclein or its fragments have been found accumulating in postmortem Alzheimer’s disease, Parkinson’s disease and Dementia with Lewy Bodies brains. Although exact mechanisms are unclear, α-synuclein is normally involved in synaptic function (9). Misfolding, mutation or post-translational modification leading to its accumulation is potentially pathogenic. Mechanisms of α-synuclein toxicity include disruption of protein trafficking, interference with the autophagy-lysosomal and mitochondrial function, thus contributing to neurodegeneration and Parkinson’s disease pathogenesis and progression (10-12). Importantly, recent studies have found that α-synuclein can spread from a donor cell to neighboring cells and thus propagate cellular damage, antagonizing the effectiveness of therapies such as transplantation of fetal or iPSC derived dopaminergic cells (13-17). While α-synuclein spreading involves multiple mechanisms including endocytosis, direct penetration, trans-synaptic transmission or through membrane receptors (11), a critical component may be autophagic failure which has been shown to promote the intercellular transfer of α-synuclein (18-20).

Multiple approaches have been explored to target α-synuclein for therapeutic development (21, 22). Proteasomal and lysosomal pathways can degrade α-synuclein *in vivo* but these functions decline both with age and with the development of Parkinson’s disease (23). There is an urgent unmet need to investigate novel approaches to enhance the clearance of α-synuclein protein. Autophagy is attractive as a therapeutic strategy because it has the potential to not only remove excessive or toxic α-synuclein species, but also degrade other damaged cellular constituents. In this regard, rapamycin and other autophagy activators have been shown to be protective in cell and animal models of neurodegenerative diseases (24). However, rapamycin may promote a starvation response that degrades both healthy and unhealthy components of the cells. Furthermore, the potential benefits of inducing autophagy initiation are dependent on optimal lysosomal function which appears to be inhibited in Parkinson’s disease. Importantly, lysosomal perturbation has been implicated in Parkinson’s disease or *parkinsonism* genetic mutations or polymorphisms, including *LRRK2, SNCA, LAMP3, GBA*, and *ATP13A2* among others (25). Variants in at least 11 genes associated with Parkinson’s disease risk are involved in or disrupt the autophagy-lysosome pathway (25), suggesting a significant impact of lysosomal perturbation in Parkinson’s disease pathogenesis.

Lysosomes are particularly important because of their high capacity and their central role in converging autophagic and endosomal activities and in degradation of macromolecules and organelles. Insufficient lysosomal turnover of the pathologic α-synuclein can result in α-synuclein accumulation (26, 27). Supporting the importance of lysosomal enzymes, Cathepsin D homozygous inactivation caused neuronal ceroid lipofuscinosis with postnatal respiratory insufficiency, status epilepticus, and death within hours to weeks after birth (28). A patient with significant loss of cathepsin D enzymatic function due to compound heterozygous missense mutations, exhibited childhood motor and visual disturbances, cerebral and cerebellar atrophy, and progressive psychomotor disability (29). In addition, cathepsin D mutations in humans, sheep, and mice resulted in neurodegeneration and accumulation of α-synuclein (26, 27), indicating that cathepsin D is essential for α-synuclein clearance. Cathepsin D levels are decreased in substantia nigra neurons of Parkinson’s disease patients compared to age-matched controls (23), and decreased cathepsin D activity has been found in postmortem brains of Parkinson’s disease patients compared to control (30). We have shown that deficiency of cathepsin D leads to α-synuclein accumulation *in vivo* despite a compensatory increase in cathepsin B (26, 27). Furthermore, overexpression of cathepsin D in cell lines and worms decreases α-synuclein aggregation and cell death (26).

Studies have identified roles of TFEB as a transcription factor that activates the expression of genes encoding lysosomal proteins (31, 32). In contrast, limited studies have explored the TFEB counterpart, ZKSCAN3, which is a transcription repressor that suppresses the expression of genes encoding lysosomal proteins (33), and thus presents an opportunity as a novel target for therapeutic development. ZKSCAN3 has been reported to promote cell proliferation and its silencing inhibited cancer cell growth while having no impact on apoptosis (33). We previously found that whole body knockout of ZKSCAN3 attenuates bacterial load in lung infection (34). Previous studies using a whole body ZKSCAN3 knockout (KO) mouse identified no overt phenotypes (35) or changes in the expression of selected autophagy genes or levels of LC3II protein in brains of 8 week old animals (35). However, in response to bacterial infection in the lung, we found significant changes of ZKSCAN3 levels (34). Furthermore, using “floxed” ZKSCAN3^f/f^ mice crossed with actin-cre transgenic mice to generate conditional ZkSCAN3 knockout mice, we have demonstrated that ZKSCAN3 disruption attenuated bacterial load when lungs were exposed to *P. aeruginosa*.

We here take two contrasting and innovative approaches testing novel strategies to enhance the capacity of the autophagy-lysosomal system. First we used our newly developed genetic model with cathepsin D transgenic overexpression in central nervous system (CNS) (36). In the second approach we used mice with CNS knockout of ZKSCAN3, which normally represses transcription of autophagy and lysosome genes (33). With these new animal models, we tested the hypothesis that *strategies that enhance lysosomal function can decrease pathogenic α-synuclein burden and neurodegeneration in vivo*. After injecting either monomeric α-synuclein, or preformed α-synuclein fibrils into the striatum of wildtype, CNS-cathepsin D overexpressing or CNS-ZKSCAN3 knockout mice, we did not find significant genotype-dependent differences in p129-α-synuclein aggregate accumulation or dopaminergic neurodegeneration. These results provide key information that neither CNS-CD overexpression nor CNS-ZKSCAN3 knockout was sufficient for neuroprotection against α-synuclein pre-formed fibrils.

## Methods

### Mice

CDfloxedstop mice were generated by GenOway. STOP cassette is flanked by loxP sites, followed by human CD cDNA. When CDfloxedstop mice are bred with *Nestin-Cre* mice (Jackson laboratory), the STOP cassette between the *loxP* sequences is deleted by Cre in Nestin expressing cells to allow CD expression (N-CDtg) in the central nervous system (CNS) (36). The genotyping primers are: PCR#1 (2240 bp): F: CATGGTAAGTAAGCTTGGGCTGCAGG; R: ACGTCAGTAGTCATAGGAACTGCGGTCG. PCR#2 (Cre-excised product: 325 bp): F: AGCCTCTGCTAACCATGTTCATGCC; R: GCGGATGGACGTGAACTTGTGC. Cre: (480 bp) F: TCGCGATTATCTTCTATATCTTCAG; R: GCTCGACCAGTTTAGTTACCC. ZKSCAN3 knockout (KO) mice were generated by breeding *Zkscan3*^*f/f*^ mice (34) with *Nestin-Cre* mice (CNS-*Zkscan3KO*). Primers for genotyping are: NDEL1: AGG CCA TGC CTT AAT GGG TGG, NDEL2: TGA TGT CAA CAG CAC TGC CTT GG; and SC1: GGT TTG GTT TTG CCT GGT GCA AAT G. All mouse experiments were done in compliance with Institutional Animal Care and Use Committee guidelines.

### Generation of pre-formed fibrils of α-synuclein (PFF)

The α-synuclein gene was cloned into pRKl72 and expressed in *E*.*coli* (13-16, 37). Bacteria grown under antibiotic selection will be harvested, homogenized and dialyzed before purification through size exclusion and ionic exchange columns. 30 mg/ml of protein in 300 µl was incubated at 37ºC for 1 week to produce fibrils and then frozen at 5 mg/ml until use. Thioflavin S and protein sedimentation assays was used to confirm the generation of insoluble fibrils. At the day of fibril transduction, 20 µl of 5 mg/ml fibrils were thawed, diluted to 1 ml with PBS, then sonicated and injected (13-16, 37).

### Stereotaxic injection

We used intrastriatal injection (coordinates are: A/P:+1.0, M/L:-2, D/V:-3.0) of PFF which has been previously shown to spread over broad brain areas and induce neurological and neurodegenerative deficits (13). PFF have been shown to seed the aggregation of endogenous mouse α-synuclein in 2-3 month old wildtype mice at 90d post single unilateral injection at 5 µg per brain into the dorsal striatum (13). α-synuclein accumulation is seen in broad brain areas at 90 and 180 d post-injection. Loss of substantia nigra tyrosine hydroxylase (TH) positive neurons is evident at 180 d post injection (13).

### Perfusion and fixation

Mice were perfused with PBS, and then 4% paraformaldehyde. The brains were removed and post-fixed in 4% PFA for 48 h, then transferred to 30% sucrose. When brains sank to the bottom of solution, they were picked out and divided into anterior and posterior halves, embedded in OCT, then stored in -80°C freezer.

### Immunohistochemistry

Each brain half was sectioned on a cryostat at into 40µm thick coronal sections and stored in cryoprotectant at -20°C. Anterior sections were placed in a 12-well plate with each well containing sections with section interval = 12, while posterior sections were placed in 6 wells of a 12-well plate with each well containing sections with section interval = 6. For TH staining, one randomly selected well of posterior sections for each subject was washed 5 times in 1X TBS to remove cryoprotectant. Sections were then incubated in 0.3% H_2_O_2_ for 20 min, washed 3 times with TBS, and blocked in TBS with 0.25% Triton X-100 (TBS-Tx) supplemented with 5% goat serum for 1 hour at 4°C. After washing 3 times with TBS, sections were incubated in 1:5000 dilution of rabbit anti-TH antibody (Millipore, # MA152, St. Louis, MO) in TBS with 5% goat serum at 4°C overnight. Sections were washed 3 times with TBS-Tx to remove unbound primary antibody then prepared using the Rabbit IgG VECTASTAIN® Elite ABC-HRP Kit (PK-6101, Vector Laboratories, Burlingame, CA). Briefly, sections were incubated in 1:500 dilution of biotinylated goat anti-rabbit secondary antibody in TBS with 5% goat serum for 1 hour at room temperature, washed 3 times with TBS-Tx, incubated in avidin–biotin substrate for 30 min, and then washed 3 times with TBS. Next, sections were incubated in 3,3’-diaminobenzidine (DAB) solution for 1-2 min (SK-4100, Vector Laboratories) and the DAB staining reaction aborted with 3 washes in ddH_2_O. Sections were mounted on glass slides and dried overnight at room temperature. Slides were then dehydrated by submersion in ddH2O for 30 seconds followed by two 3 minute incubations in increasing concentrations of ethanol (70%, 95%, and 100%). Slides were then washed for 5 minutes 3 times in HistoClear (50-899-90147, Fisher Scientific) and briefly dried before the addition of Permount mounting media (SP15-100, Fisher Scientific) and coverslips. Slides were laid flat on the bench and mounting media cured overnight before storage, imaging, and quantification.

For p-α-synuclein staining, we used mouse anti-mouse p-α-synuclein antibody (anti-Phospho (Ser129)-α-Synuclein; Abcam EP1536Y), and Alexa Fluor 488 goat anti-rabbit IgG for p-Syn. Sections were mounted on Fischer SuperFrost Plus microscope slides partially submerged in TBS using a paintbrush. ProLong Diamond Antifade mountant with DAPI (Fisher: P36971) was applied and a coverslip was placed over the slide.

### Unbiased Stereological analysis

On each slide, TH+ neurons in the substantia nigra (SN) were quantified by unbiased stereology using an Olympus BX51 microscope and Stereo Investigator software (version 2020; MBF Bioscience, Williston, VT) Optical Fractionator workflow. Sections containing SN were identified by the pattern of TH staining and visual comparison of morphological features to representative coronal sections of mouse brain using the Allen Brain Atlas (https://mouse.brain-map.org/static/atlas). On 4 sections containing SN, contours defining the regions of interest (ipsilateral and contralateral to the injection site) at 4X magnification were placed. Average mounted thickness of similarly prepared sections was previously determined to be approximately 30µm (unpublished data) and optical dissector height (on the z-axis) set to 20µm. For systematic random sampling, a 100µm x 100µm grid was placed over the region of interest and 50µm x 50µm counting frame established. TH+ neurons within the counting frame and optical dissector were counted at 40X magnification and population estimates for TH+ neurons in each hemisphere generated by the software.

### Statistics

We used two-way, and three-way ANOVA analyses, followed by Fisher’s LSD as appropriate. The statistics used are indicated in the results and figure legends.

## Results

### Neither central nervous system (CNS) cathepsin D overexpression (CNS-CDtg) nor CNS-ZKSCAN3 knockout (CNS-ZkscanKO) in the brain protect against dopaminergic neurodegeneration induced by intrastriatal pre-formed α-synuclein fibrils (PFF) injection

Previous studies demonstrated that intracranial injection of α-synuclein monomers does not cause neurodegeneration (38). At 4 months post-injection, there was no difference due to genotype or treatment in the relative reduction of TH+ neurons in the hemisphere ipsilateral to the injection site in Control and CNS-CDtg mice that had been injected with monomeric α-synuclein versus those injected with PFFs [**Figure 1A** (4month) and **Table 2** (THStereology); F_INT_(1,28) = 0.29, p = 0.5933; F_GENO_(1,28) = 0.10, p = 0.7510; F_Tx_(1,28) = 0.38, p = 0.5430].

**Table 1.** The numbers of mice used. We injected either α-synuclein monomer or pre-formed fibrils (PFF) into the striatum (all male, at 2-3 months of age) and perfused mice either 4 months or 6 months post-injection.

| Genotype | 4 months PFF | 4 months monomer | 6 months PFF | 6 months monomer |
| --- | --- | --- | --- | --- |
| Controls<br>(*Nestin-cre*, *CDfloxedstop*, *Zkscan3<sup>fl/+</sup>*, and *Zkscan3<sup>fl/fl</sup>*) | 17 | 2 | 20 | 5 |
| CD overexpression in the brain<br>(CNS-*CDtg*) | 13 | 2 | 17 | 3 |
| ZKSCAN3 knockout in the brain<br>(CNS-*Zkscan3*KO) |  |  | 9 | 4 |

**Table 2.** Quantification of dopaminergic neuron TH+ staining in the substantia nigra after monomer or PFF injection.

|  | Monomer |  |  |  | PFF |  |  |  |
| --- | --- | --- | --- | --- | --- | --- | --- | --- |
|  |  | Contralateral Population Estimate | Ipsilateral Population Estimate | Ipsilateral % of Contralateral |  | Contralateral Population Estimate | Ipsilateral Population Estimate | Ipsilateral % of Contralateral |
| **4 Months** | **n** | **Mean ± SD** | **Mean ± SD** | **Mean ± SD** | **n** | **Mean ± SD** | **Mean ± SD** | **Mean ± SD** |
| **Control** | 2 | 6948 ± 407 | 5346 ± 586 | 76.83 ± 3.92 | 16 | 8127 ± 1471 | 6473 ± 1848 | 78.37 ± 10.84 |
| **CNS-CDtg** | 2 | 6678 ± 891 | 5688 ± 967 | 84.96 ± 3.15 | 12 | 8757 ± 2225 | 6951 ± 2361 | 78.91 ± 16.67 |
| **6 Months** |  |  |  |  |  |  |  |  |
| **Control** | 5 | 9086 ± 1829 | 8914 ± 1845 | 98.37 ± 6.38 | 17 | 7376 ± 1788 | 5315 ± 1424 | 72.94 ± 13.40 |
| **CNS-CDtg** | 3 | 10524 ± 2453 | 9528 ± 1952 | 90.98 ± 6.11 | 14 | 7791 ± 2080 | 5863 ± 1382 | 76.62 ± 10.22 |
| **CNS-ZkscanKO** | 3 | 7332 ± 4202 | 6888 ± 3420 | 96.39 ± 13.81 | 9 | 7036 ± 2152 | 4648 ± 1676 | 67.07 ± 13.58 |

**Figure 1.**
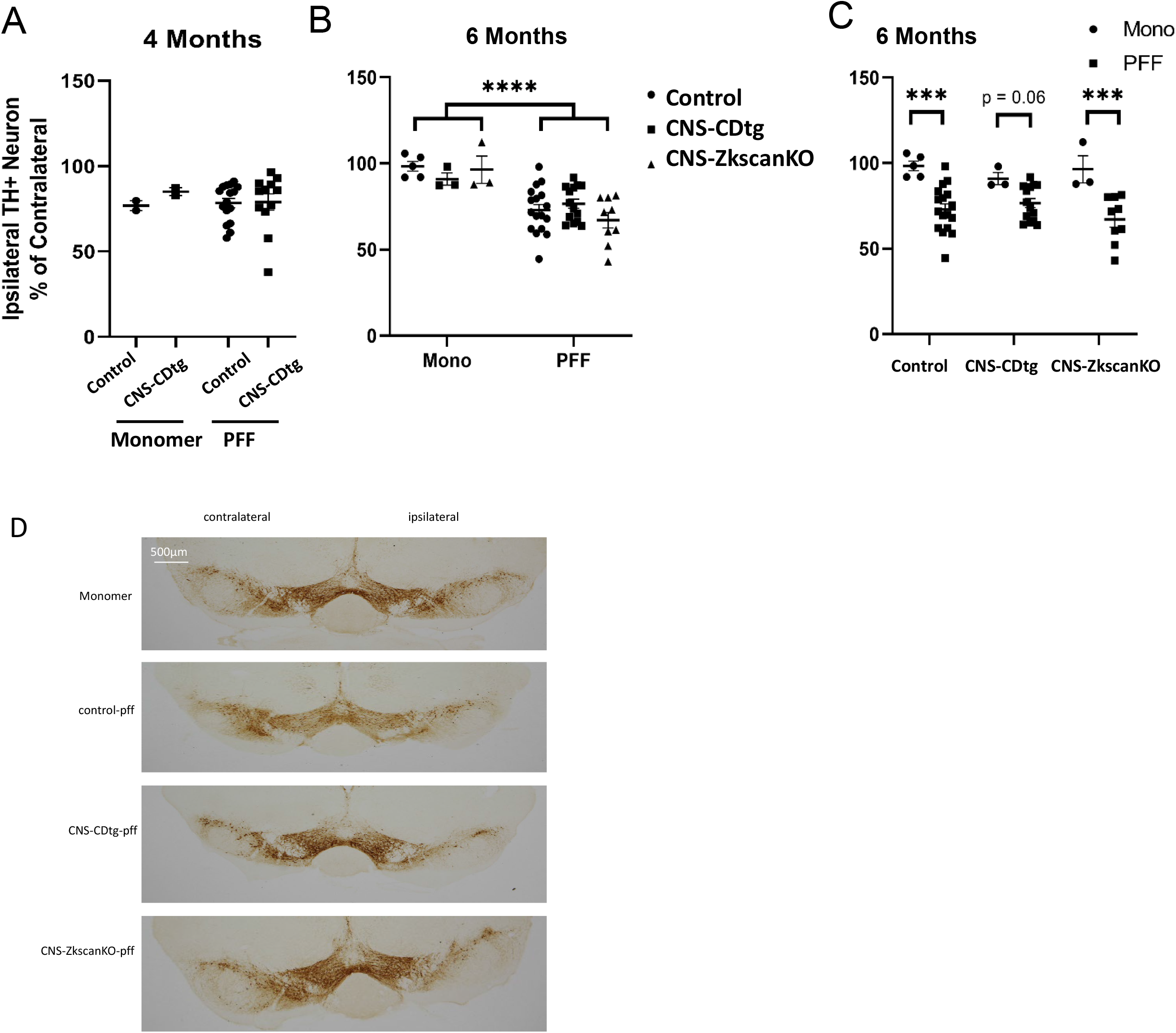
Neither cathepsin D overexpression nor ZKSCAN3 knockout in the central nervous system (CNS) (CNS-CDtg and CNS-ZkscanKO) protect against dopaminergic neurodegeneration induced by preformed α-synuclein fibrils (PFF). **A)** Quantification of TH-positive neurons in the substantia nigra 4 months following unilateral striatal injections of either monomeric or PFF of α-synuclein. Control and CNS-CDtg mice received unilateral striatal injections of either monomer or PFF α-synuclein. After 4 months, the mice were perfused, sections immunostained, and unbiased stereology of TH+ neurons performed by an investigator blinded to experimental conditions. No significant differences were detected between control and CNS-CDtg mice 4 months following monomer or PFF injection. To account for variability in staining intensity between animals, the population estimate of the ipsilateral hemisphere is expressed as a percent of the population estimate of the contralateral hemisphere with mean and S.E.M. indicated. Data were analyzed by two-way ANOVA. **B)** Quantification of TH-positive neurons in the substantia nigra 6 months following unilateral striatal injections of either monomeric or PFF of α-synuclein. Control, CNS-CDtg, and CNS-ZkscanKO mice received unilateral striatal injections of either monomer or PFF α-synuclein. After 6 months, the mice were perfused, immunostaining was performed, and unbiased stereology of TH+ neurons performed by an investigator blinded to experimental conditions. To account for variability in staining intensity between animals, the population estimate of the ipsilateral hemisphere is expressed as a percent of the population estimate of the contralateral hemisphere with mean and S.E.M. indicated. PFF-injected mice have a greater reduction of TH+ neurons ipsilateral to the injection site than those injected with monomeric α-synuclein. **C)** Data in panel B was plotted and analyzed with each genotype. PFF-injected control and CNS-ZkscanKO mice have 25% and 29% less ipsilateral TH+ neurons relative to those that were monomer-injected. PFF-injected CNS-CDtg mice demonstrated 14% fewer ipsilateral TH+ neurons than monomer-injected CNS-CDtg mice, however, this reduction failed to meet the threshold for significance. ***p < 0.001, ****p < 0.0001. **D)** Representative TH immunostaining from 6 months post injection with monomer or PFF in the 3 genotypes.

At 6 months post-injection of monomeric α-synuclein, TH+ neurons in the hemisphere ipsilateral to injection compared to the TH+ neurons in the uninjected hemisphere were 98.4%, 91.0%, and 96.4%, in Control, CNS-CDtg, and CNS-ZkscanKO mice respectively (**Table 2** (THStereology)). At 6 months post-injection of PFFs, TH+ neurons in the hemisphere ipsilateral to injection compared to the TH+ neurons in the uninjected hemisphere were 72.9%, 76.6%, and 67.1%, in Control, CNS-CDtg, and CNS-ZkscanKO mice (**Table 2** (THStereology)). Two-way ANOVA confirmed the expected difference in relative ipsilateral TH+ neuron reduction between mice treated with monomeric α-synuclein and those treated with PFFs [F_Tx_(1,45) = 30.67, p < 0.0001], but failed to identify a specific effect of genotype [F_GENO_(2,45) = 0.32, p = 0.7308], or an interaction between effects of genotype and treatment [F_INT_(2,45) = 1.06, p = 0.3564] (**Figure 1B** (6month). Comparisons within each genotype found that PFF treatment resulted in greater TH+ neuron loss in Control and CNS-ZkscanKO mice [t_WT_(45) = 4.22, p = 0.0001, q = 0.0004; t_ZKKO_(45) = 3.71, p = 0.0006, q = 0.0009]; but the trend toward greater neuron loss was not significant in CNS-CDtg mice [t_CDTG_(45) = 1.90, p = 0.0634] (**Figure 1C-D** (6month) and **Table 2** (THStereology). Comparisons of CNS-CDtg mice treated with PFF at 4 months versus 6 months post-injection found no difference in the relative reduction of TH+ neurons [t(24) = 0.43, p = 0.6724]. This result suggests that CNS-CDtg may be protective of substantia nigra dopaminergic cell death at 6 months post-injection.

### Neither CNS-CDtg nor CNS-ZkscanKO in the brain protect against pS129 α-Synuclein aggregates in the ipsilateral side of PFF injection in the substantia nigra

Control and CNS-CDTG mice injected with PFFs demonstrated increased pS129 α-synuclein positive aggregates relative to those injected with monomeric α-synuclein at 4 months post-injection, irrespective of genotype [**Figure 2A** (4month) and **Table 3** (aSYN); F_INT_(1,26) = 0.04, p = 0.8461; F_GENO_(1,26) = 0.04, p = 0.8461; F_Tx_(1,26) = 6.80, p = 0.0149]. In mice injected with PFFs, there was no difference in the number, area, or mean size of pS129 α-synuclein puncta [**Table 3** (aSYN); t_#_(25) = 0.64, p = 0.5275); t_area_(25) = 1.39, p= 0.1763; t_size_(25) = 1.45, p = 0.1594] 4 months after injection.

**Table 3.**
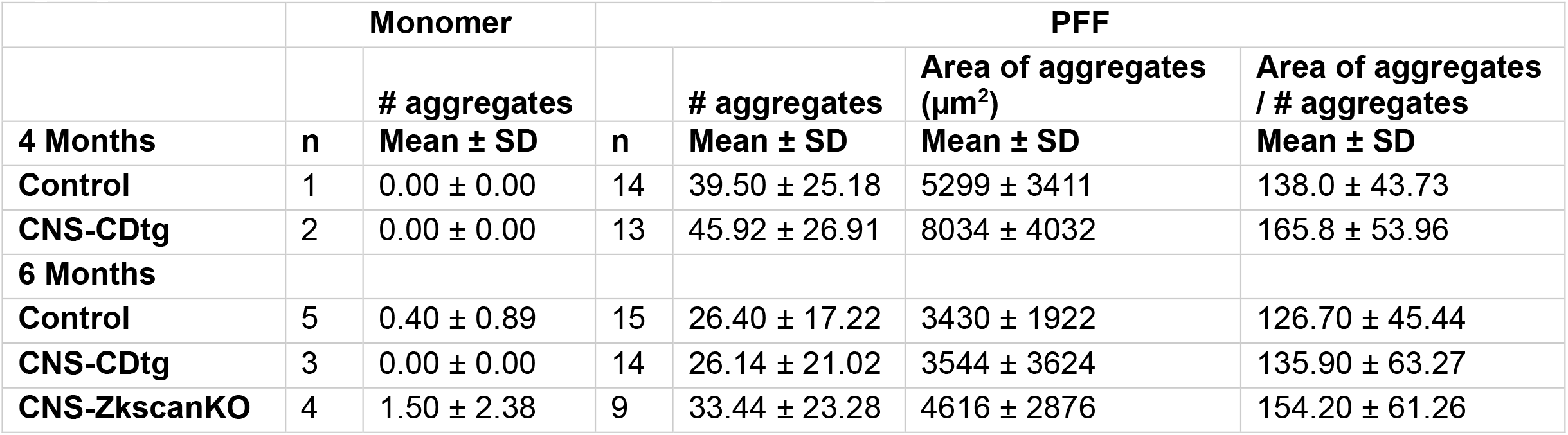
pS129 α-synuclein aggregates in control, CNS-CDtg, and CNS-ZkscanKO mice in the substantia nigra ipsilateral to either monomeric or PFF α-synuclein injection.

**Fig. 2.**
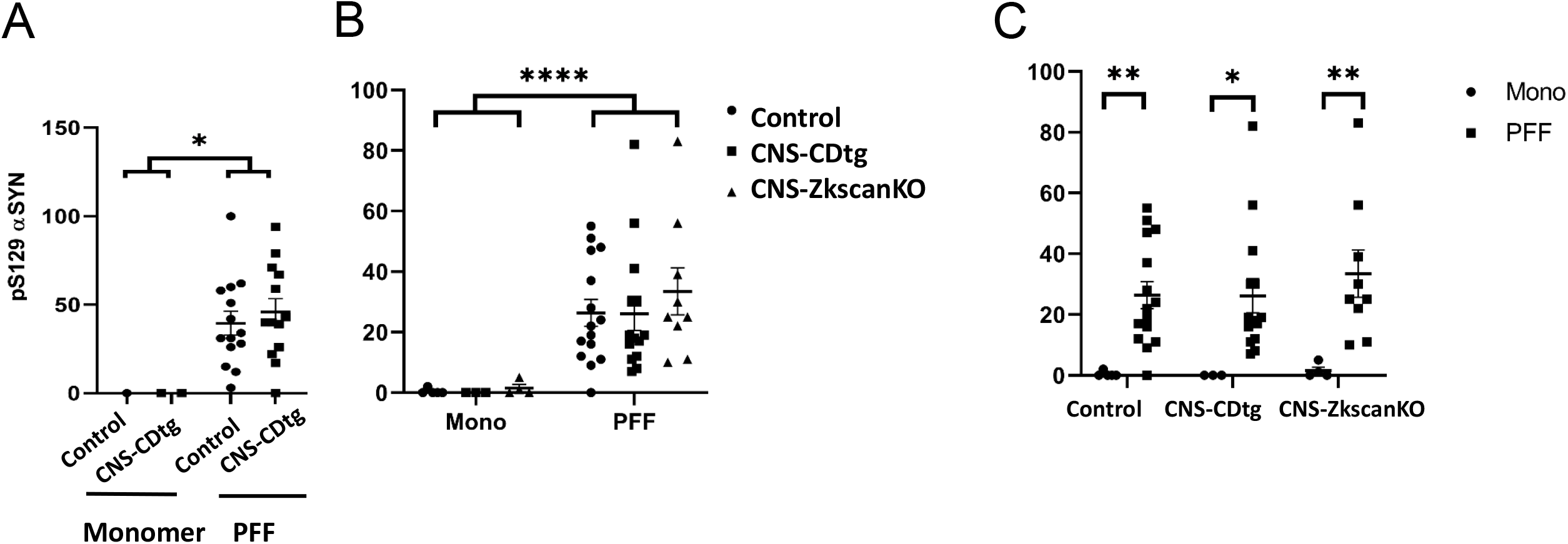
p-syn quantification in substantia nigra (SNr). A) Significant increase of the number of pS129 α-synuclein aggregates ipsilateral to the injection site after 4 months of PFF injection compared to the monomer. Data were analyzed by two-way ANOVA and Fisher LSD tests. *p < 0.05. B) Significant increase of the number of pS129 α-synuclein aggregates ipsilateral to the injection site after 6 months of PFF injection compared to the monomer. Data were analyzed by two-way ANOVA. ****p < 0.0001. C) Increased pS129 α-synuclein aggregates in PFF treated mice were evident in each genotype. Data were analyzed with two-way ANOVA and Fisher LSD tests. *p < 0.05, **p <0.01. Data = mean ± S.E.M.

Control, CNS-CDTG, and CNS-ZkscanKO mice treated with PFFs had increased pS129 α-synuclein aggregates relative to mice treated with monomeric α-synuclein at 6 months post-injection [**Figure 2B** (6month-aSYN) and **Table 3** (aSYN); F_INT_(2,44) = 0.10, p = 0.8461; F_GENO_(2,44) = 0.21, p = 0.8113; F_Tx_(1,44) = 21.15, p < 0.0001]. This effect was significant in each genotype [**Figure 2C** (6month-aSYN) and Table 3 (aSYN); t_WT_(44) = 2.80, p = 0.0076; t_CDTG_(44) = 2.28, p = 0.0273; t_ZKKO_(44) = 2.95, p = 0.0050]. There was no difference between groups treated with PFFs in aggregate area or mean size 6 months post-infection (**Table 3** (aSYN)). These results suggest that CNS-CDtg is still permitting p-α-synuclein accumulation at 4 months but less so at 6 months, indicating a delay from the p-α-synuclein accumulation and its clearance.

### Comparison between controland CNS-CDtg mice 4 months and 6 months post-injection in the substantia nigra

Comparing the number of pS129 α-synuclein positive aggregates in CNS-CDtg and control mice treated with either monomeric or PFF α-synuclein at 4 months and 6 months post-injection found that the type of α-synuclein (monomer vs PFF) accounted for 19.6% of the total variation between groups [**Figure 3A left** (4mo v 6mo); F_Tx_(1,59) = 17.94, p < 0.0001]. Similarly, mice injected with PFFs had a greater relative reduction of TH+ neurons ipsilateral to the injection [**Figure 3B left** (4mo v 6mo); F_Tx_(1,63) = 7.34, p = 0.0087] and an interaction between treatment and months post-injection was identified [F_Tx x months_(1,63) = 4.66, p = 0.0348]. No specific effects of genotype or other interactions between genotype, treatment, and months post-injection in the relative reduction of TH+ neurons were identified.

**Fig. 3.**
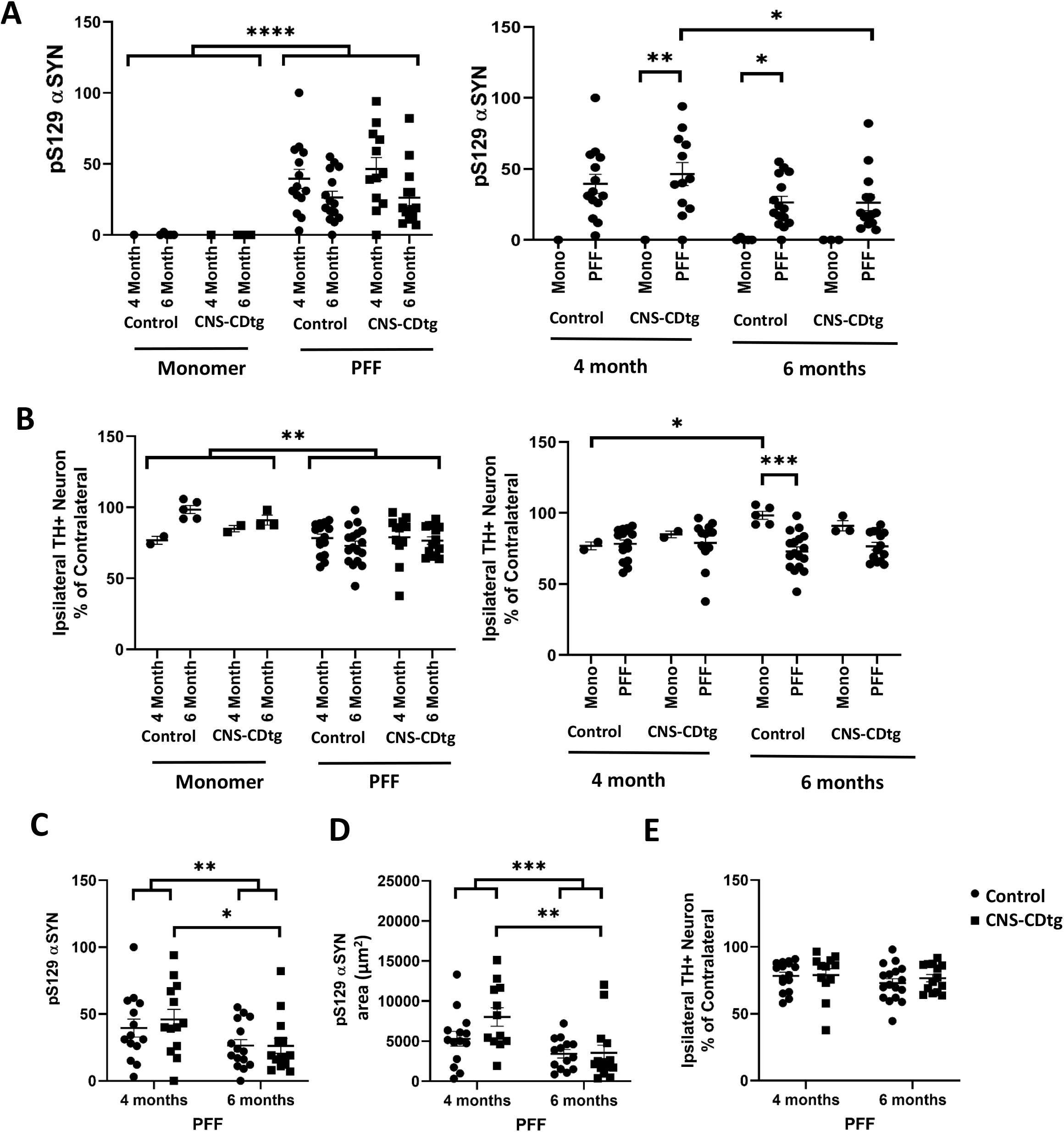
Comparison of pS129 α-synuclein aggregates and TH+ neurons in control and CNS-CDtg mice 4 months and 6 months after unilateral striatal injections of either monomer or PFF. A) Control and CNS-CDtg mice treated with PFF demonstrated a higher number of pS129 α-synuclein aggregates at both 4 and 6 months post-injection relative to monomer injected mice (left). Comparing the difference between PFF and monomer injected mice 4 and 6 months post injection, CNS-CDtg mice have more aggregates 4 months after treatment with PFF, while PFF treated control mice have more aggregates than monomer treated 6 months after injection (right). CNS-CDtg mice also have fewer pS129 α-synuclein aggregates at 6 months compared to 4 months (right). Data are shown as the number of pS129 α-synuclein puncta ipsilateral to the injection site with mean and S.E.M indicated. B) Control and CNS-CDtg mice treated with PFF had fewer TH+ neurons than monomer treated at 4 and 6 months post-injection (left). Control mice injected with monomeric α-synuclein had less relative reduction of TH+ neurons at 6 months than at 4 months, and PFF injected control mice had greater relative reduction of TH+ neurons than monomer treated at 6 months post-injection (right). The population estimate of TH+ neurons in the ipsilateral hemisphere is expressed as a percent of the population estimate of the contralateral hemisphere with mean and S.E.M. indicated. C-D) Control and CNS-CDtg mice treated with PFF had fewer pS129 α-synuclein aggregates and the area of aggregates present was smaller at 6 months relative to at 4 months post-injection, The reduction in number and area of α-synuclein aggregates in CNS-CDtg mice is significantly greater at 6 months than 4 months post-injection. E) There is no difference in the relative reduction of TH+ neurons following PFF treatment in control or CNS-CDtg mice between time points. Data in panels A and B were analyzed by three way ANOVA and Fisher’s LSD tests, and data in panels C-E were analyzed by two-way ANOVA and Fisher’s LSD tests. *p < 0.05, **p < 0.01, ***p < 0.001, ****p < 0.0001

At 6 months post-injection, a greater number of pS129 α-synuclein aggregates were detected in PFF treated control mice compared to monomer treated [**Figure 3A right** (4mo v 6mo); t(59) = 2.36, p = 0.0215], and TH+ neuron loss was also greater in PFF versus monomer treated control mice at 6 months [**Figure 3B right** (4mo v 6mo); t(17) = 4.11, p = 0.0001]. Differences between monomer and PFF treatment on the number of pS129 α-synuclein puncta in CNS-CDtg mice were evident at 4 months post-injection but this effect failed to reach the threshold for significance at 6 months [**Figure 3A right** (4mo v 6mo); t_4month_(59) = 2.84, p = 0.0063; t_6month_(14) = 1.85, p = 0.0681]. Interestingly, in CNS-CDtg mice, the number of pS129 α-synuclein aggregates at 6 months was 45% less than the number of puncta detected at 4 months. [**Figure 3A right** (4mo v 6mo); t(59) = 2.41, p = 0.0192]; although there was no corresponding difference in the relative reduction of TH+ neurons [t(24) = 0.43, p = 0.6724]. In monomer injected control mice, ipsilateral TH+ neuron levels were higher at 6 months relative to levels at 4 months [**Figure 3B right** (4mo v 6mo) t(63) = 2.12, p = 0.0381].

In PFF treated control and CNS-CDtg mice, both the number and area of pS129 α-synuclein aggregates were decreased at 6 months relative to measures taken 4 months post-injection [**Figure 3 C-D** (4mo v 6mo); F_#Puncta_(1,52) = 7.32, p = 0.0092; F_area_(1,50) = 12.37, p = 0.0009], although the mean size of α-synuclein aggregates (Area of aggregates / # aggregates, **Table 3**) were not different for either genotype at either time point [F_size_(1,50) = 2.113, p = 0.1523]. Relative reductions of ipsilateral TH+ neurons were also not different between PFF treated control and CNS-CDtg mice at either time point [**Figure 3E** (4mo v 6mo); F(1,55) = 1.31, p = 0.2570]. Interestingly, the reduction of p129 α-synuclein puncta number and area in PFF treated mice at 6 months appear to be driven by differences in CNS-CDtg mice [**Figure 3C-D** (4mo v 6mo); t_#Puncta_(52) = 2.26, p = 0.0280; t_area_(50) = 3.44, p = 0.0012].

### pS129 α-Synuclein aggregates in the cortex, striatum and amygdala

We also scored the pS129 α-Synuclein aggregates in the cortex, striatum and amygdala by investigators blind to genotype and treatment. There were no pS129 α-Synuclein aggregates in monomer injected brains. PFF injected brains were scored with 1 as >5 aggregates per section but not extensive, 2 as many aggregates but not as clustered, and 3 and many intense aggregates and clustered (**Fig.4**). There was no difference in any of the regions between control and CNS-CDtg mice at 4 months (**Fig.4A**), and no difference between 4 and 6 months for control and CNS-CDtg brains (**Fig.4B**). However, there were less aggregates in the cortex and striatum of CNS-ZkscanKO brains compared to WT and CDTG at 6 months post injection (**Fig.4C-D**). This result suggest that CNS-Zkscan KO may be effective for p-α-synuclein clearance in regions where its burden is less severe than substantia nigra.

**Fig. 4.**
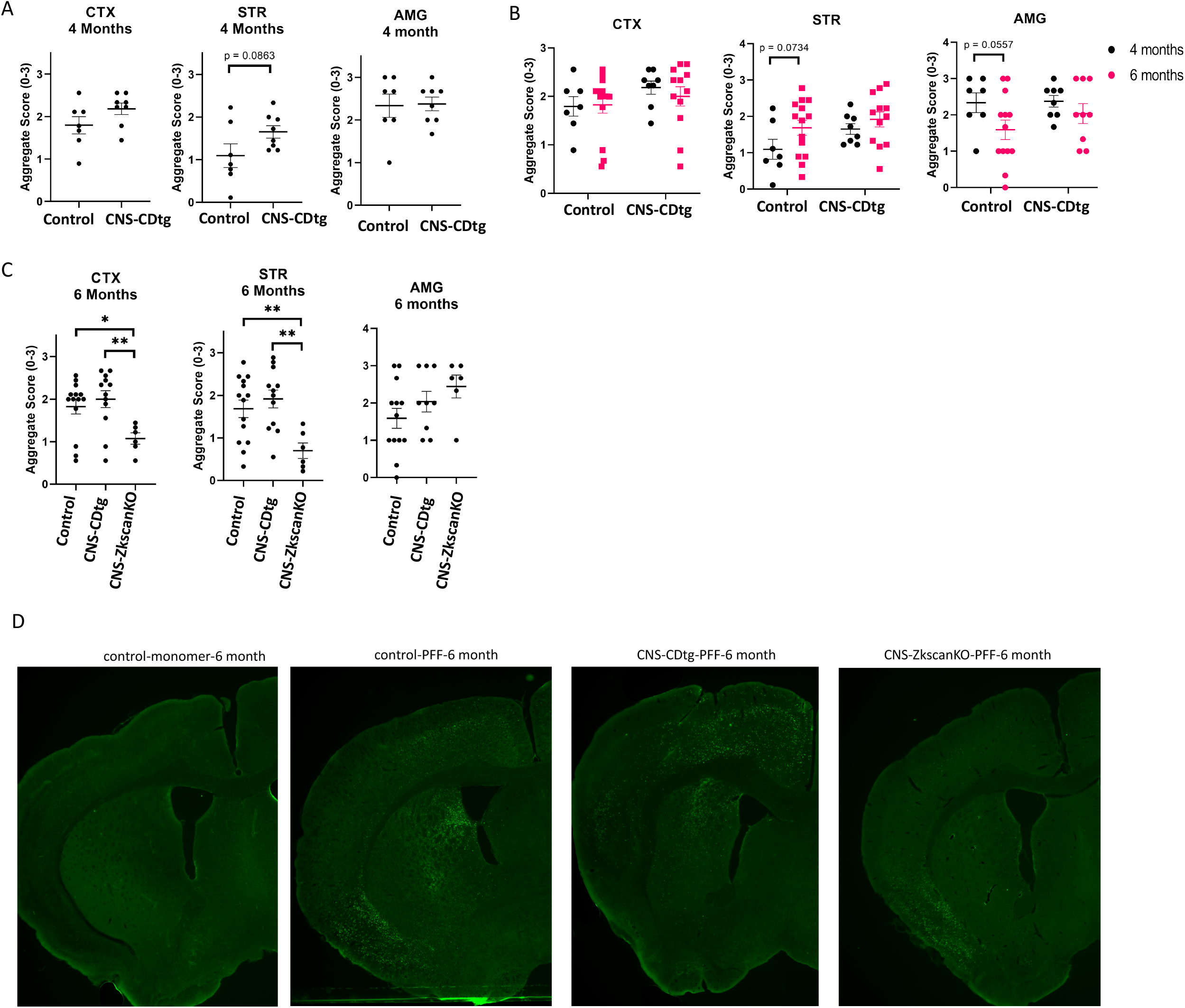
p-syn quantification in cortex, amygdala and striatum. Scoring was performed by investigators blind to genotype or treatment. There was no difference in any regions between control and CNS-CDtg mice at 4 months (A), there was also no change from 4 to 6 months in control and CNS-CDtg mice (B). In contrast, there was a decrease of aggregates in CNS-ZkscanKO mice in the cortex and striatum (C). Representative p-syn immunostaining images are shown in (D). Two-way ANOVA was performed followed by Fisher’s LSD. *p < 0.05, **p < 0.01.

## Discussion

α-synuclein aggregates can spread from a donor cell to neighboring cells and thus propagate cellular damage, an observation that may underlie the pathological progression of neurodegenerative diseases. We and others previously have shown that insufficient lysosomal function due to genetic mutations or targeted disruption of cathepsin D can cause α-synuclein accumulation (26, 27). We have also shown that knockout of lysosomal transcription suppressor ZKSCAN3 elevates key autophagy gene expression in postnatal day 10 lung and attenuates bacterial accumulation when injected to the lung (34). We here investigated whether overexpression of cathepsin D or knockout transcription suppressor of lysosomal biogenesis ZKSCAN3 in the central nervous system can attenuate α-synuclein prorogation and dopaminergic cell death after injection of pre-formed fibrils. To this end, we did not find significant genotype dependent differences. Our results indicate that enhancing cathepsin D in the CNS is not sufficient to attenuate p129-α-synuclein aggregate accumulation or dopaminergic neurodegeneration. Neither did ZKSCAN3 KO in the CNS significantly impact these parameters.

There are some modest differences observed. For example, p129-α-synuclein aggregation was significant at 4 month post-PFF injection in CNS-CDtg mice compared to post-monomer injection, but not wildtype mice, while at 6 month post-PFF injection p129-α-synuclein aggregation was significant in the wildtype but not CNS-CDtg mice compared to post-monomer injection. The loss of dopaminergic neurons 6 months post-PFF injection is significant in wildtype mice but not CNS-CDtg mice. One potential explanation is that there are more dopaminergic neurons in monomer injected wildtype mice 6 month post-injection compared to 4 months. We performed this study based on prior studies that monomers do not induce cell death or p129-α-synuclein aggregation. As we found that it is true that there was no p129-α-synuclein aggregation in response to monomer injection. However, there was a decrease of ipsilateral TH+ neurons in these 2 mice we used. Nonetheless, the observation that 6 months post PFF there was a significant decrease of TH+ neurons compared to monomer injected in wildtype mice but not CNS-CDtg mice brings up the possibility that wildtype mice may exhibit progressively higher levels of cell loss after prolonged post-PFF injection, while CNS-CDtg may not exhibit as severe cell loss. Furthermore, CNS-Zkscan KO may be effective p-α-synuclein clearance in regions where its burden is less severe than substantia nigra, such as cortex and striatum. Future studies will have to further test these hypotheses.

